# Axon guidance modalities in CNS regeneration revealed by quantitative proteomic analysis

**DOI:** 10.1101/2021.03.26.437129

**Authors:** Noemie Vilallongue, Julia Schaeffer, Anne Marie Hesse, Céline Delpech, Antoine Paccard, Yohan Couté, Stephane Belin, Homaira Nawabi

## Abstract

Long-distance regeneration of the central nervous system (CNS) has been achieved from the eye to the brain through activation of neuronal molecular pathways or pharmacological approaches. Unexpectedly, most of the regenerative fibers display guidance defects, which prevents reinnervation and further functional recovery. Therefore, characterizing the mature neuronal environment is essential to understand the adult axonal guidance in order to complete the circuit reconstruction. To this end, we used mass spectrometry to characterize the proteomes of major nuclei of the adult visual system: suprachiasmatic nucleus (SCN), ventral and dorsal lateral geniculate nucleus (vLGN, dLGN) and superior colliculus (SCol)), as well as the optic chiasm. These analyses revealed the presence of guidance molecules and guidance-associated factors in the adult visual targets. Moreover, by performing bilateral optic nerve crush, we showed that the expression of some proteins was significantly modulated by the injury in the visual targets, even in the ones most distal to the lesion site. On another hand, we found that the expression of guidance molecules was not modified upon injury. This implies that these molecules may possibly interfere with the reinnervation of the brain targets. Together, our results provides an extensive characterization of the molecular environment in intact and injured conditions. These findings open new ways to correct regenerating axon guidance notably by manipulating the expression of the corresponding guidance receptors in the nervous system.

## INTRODUCTION

In adult mammals, neurons of the central nervous system (CNS) are unable to regenerate following a lesion. Therefore, any damage to the CNS leads to permanent cognitive and motor disabilities in patients. This failure of regeneration has two main components: the inhibitory environment of the lesion site and the intrinsic properties of neurons themselves (He and Jin, 2016). Indeed, there is a progressive loss of regenerative capabilities of the CNS over the course of development (Park et al., 2008), and this decline is exacerbated after axon injury (Belin et al., 2015). Over the past few years, combined genetic manipulations and pharmacological approaches have led to robust CNS regeneration. In these models, axons are able to grow over long distances, for example up to several centimeters from the eye ball to the brain in the mouse visual system (Belin et al., 2015; Lim et al., 2016; Lu et al., 2020). However, even if some models result in partial functional recovery, there is a lack of appropriate guidance of regenerative fibers to their functional target, which explains the absence of full functional recovery (Crair and Mason, 2016; Pernet and Schwab, 2014).

Regenerative RGC display strong guidance defects already in the optic nerve, with a abnormally high number of U-turns or branches (Pernet et al., 2013b, 2013a). In long-regeneration models, the vast majority of regenerative axons are observed to get lost in the optic chiasm, fail to follow the optic tract into the brain, or even grow into the contralateral optic nerve (Belin et al., 2015; Luo et al., 2013). Furthermore, out of the axons that reach the optic tract, very few axons are actually detected in the brain targets while many clearly avoid them (Lim et al., 2016). These lost axons might jeopardize functional recovery, with several hypotheses arising: the number of regenerative axons reaching their proper target might be insufficient to reestablish functional recovery; lost axons might form aberrant connections that might impair further the formation of a proper neuronal circuit. The latter hypothesis may explain the behavioral phenotype observed in Sox11-overexpression regeneration paradigm (Wang et al., 2015): while Sox11 overexpression induces robust regrowth of corticospinal (CST) axons after spinal cord injury, the animals performed less well in forelimb function compared to controls. Therefore the burning question is whether mature axons are primed to be guided?

During embryonic development, thousands of neurons project their axons over long distances to reach their functional targets. Axon navigation is oriented by their response to guidance cues at critical choice points (Bellon and Mann, 2018; Stoeckli, 2018). These guidance molecules, either attractive or repulsive, comprise soluble molecules that can act over long distances, such as secreted Semaphorins, Slits and Netrins, and transmembrane proteins acting locally, such as Ephrins. Understanding guidance activity of a single molecule is not as straightforward as originally thought. Indeed, it is becoming clearer that guidance molecules activity is finely modulated by a number of co-factors: identity of the responding neuronal population (Nédelec et al., 2012), crosstalk of other cues or receptors (Charoy et al., 2012; Delloye-Bourgeois et al., 2015; Kuwajima et al., 2012), substrate stiffness (Koser et al., 2016), neuronal activity (Nicol et al., 2007; Plazas et al., 2013). In addition, over the past few years others molecules have been described to be involved in axon guidance such as morphogens (Shh, Wnt), growth factors (TGFb, BMP) (Yam and Charron, 2013) or adhesion molecules (L1CAM, NrCAM) (Pollerberg et al., 2013).

In the context of adult regeneration in the CNS, many unknowns of axon guidance regulation remain. Thus, characterizing and controlling the neuronal environment is the critical first step towards achieving correct guidance of regenerative axons to their proper target and obtain functional recovery. So far, most studies have focused on characterizing the lesion site, which is considered to be the main barrier for axon growth (Bradbury and Burnside, 2019; Silver and Miller, 2004). Yet, long-distance models of CNS regeneration prove that what needs to be overcome is not the growth-inhibitory nature of the lesioned CNS, but rather the failure of reinnervation of proper brain targets. In this work, we sougth to characterize molecular events beyond the lesion site (located right behind the eye ball) in distal regions of the visual system. We used mass spectrometry (MS)-based proteomics to analyze the protein content of the functional nuclei of the mouse adult visual system innervated by the retina ganglion cells (RGC) axons that form the optic nerve. We also analysed the optic chiasm, a major guidance choice-point of developing RGC axons that bears many guidance defects when it comes to regeneration (Luo et al., 2013; Belin et al., 2015; Crair and Mason, 2016). Our analysis revealed a number of guidance and guidance-associated factors, notably adhesion molecules and extracellular matrix components. Furthermore, we performed bilateral optic nerve crush to trigger denervation of visual targets and mimic the environment that regenerative axon have to face. Quantitative proteomic comparison with an intact brain revealed significant changes in terms of protein expression in the damaged system. Interestingly, we found the expression of guidance cues and receptors remain steady upon injury, suggesting that the adult brain has an intrinsic guidance signature that may affect navigation of regenerating axons and more generally the connectivity of the injured circuit.

## RESULTS

### MS-based proteomic analysis of visual targets in the adult brain

The development of the visual system, which is composed of the retina, optic nerve, optic tracts and functional nuclei in the brain, has been thoroughly studied during development (Hong and Chen, 2011; Seabrook et al., 2017). Within the retina, the RGC population is the only one to project its axon to the brain via the optic nerve. Most of the topographic maps are very well established (Huberman et al., 2008; Seabrook et al., 2017; Wienbar and Schwartz, 2018). Guidance molecules that control RGC projections have been identified, in particular for the process of midline crossing at the optic chiasm (Erskine and Herrera, 2014). Regarding the functional targets, several recent transcriptomics studies have unraveled gene expression regulation during development and in adult (Kalish et al., 2018; Masullo et al., 2019; Wang et al., 2016; Wen et al., 2020). However, it is now clear that protein and mRNA levels do not correlate (Komili and Silver, 2008; Liu et al., 2016; Vogel and Marcotte, 2012), meaning that the transcriptomic profiling of a particular tissue or condition does not necessarily reflect the protein expression. Therefore, we used mass spectrometry-based proteomics to characterize the protein composition of the mature brain regions targeted by RGC axons (**Figure 1A**). To this end, we injected fluorescently-labelled cholera toxin B (CTB) in mouse eyes and microdissected the selected target regions in the brain: suprachiasmatic nucleus (SCN), dorsal and ventral lateral geniculate nucleus (dLGN and vLGN) and superior colliculus (SCol) (**Figure 1B-C**). Since the optic chiasm is a critical intermediate target where the majority of RGC axons get lost during regeneration, we also microdissected adult optic chiasm to decipher particular extracellular signals that may influence guidance in adult (**Figure 1B-C**).

**Figure 1:**
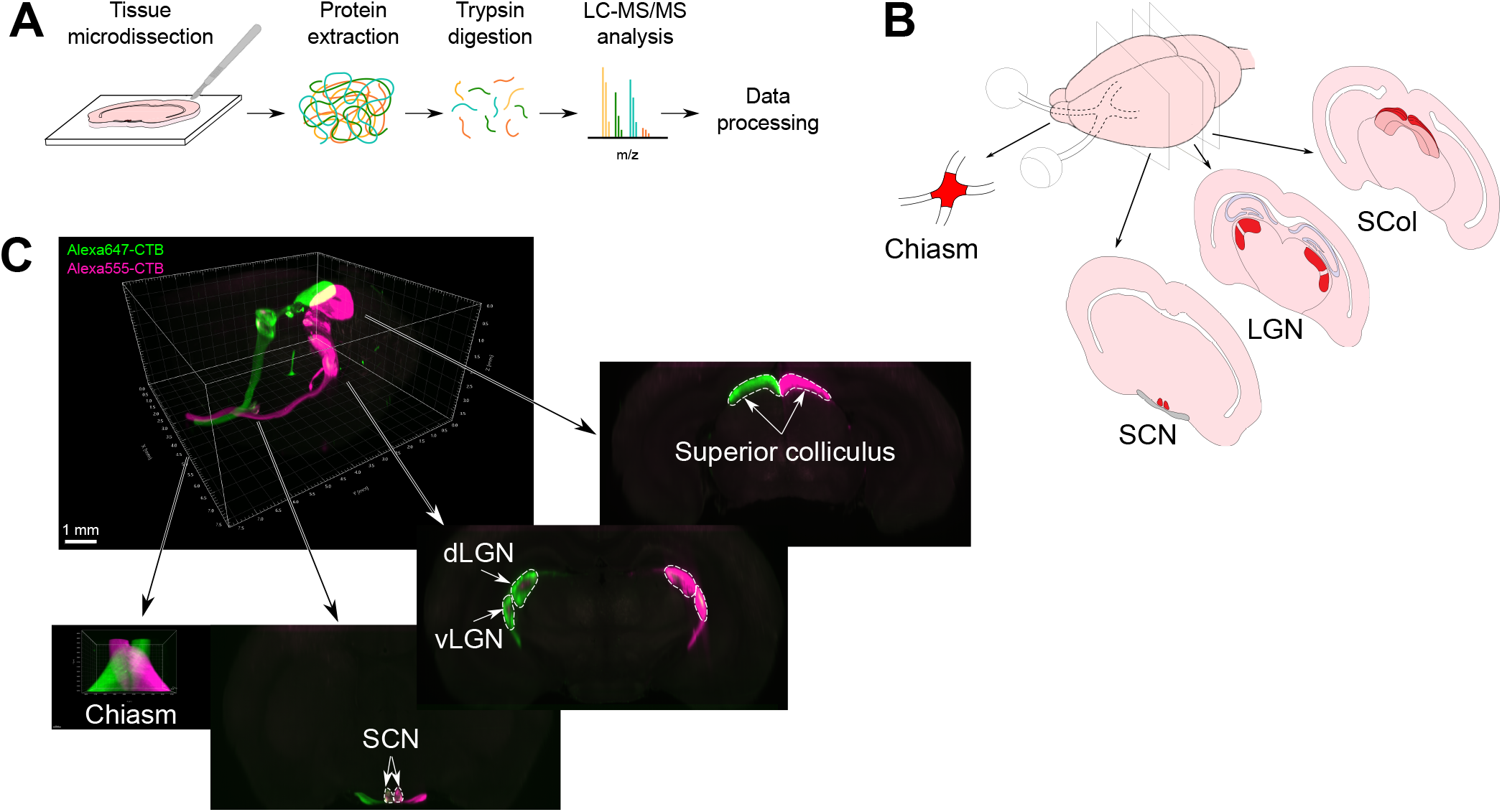
Proteomics analysis of adult visual targets. **(A)** Experimental design. Proteins were extracted from microdissected tissues and analysed by LC-MS/MS after trypsin digestion. **(B)** Diagram of the mouse visual targets used for the study. Primary targets of RGC axons (indicated in red) and the optic chiasm were collected for the MS analysis. **(C)** 3D transparized brain imaged using a lightsheet confocal microscopy. The two eyes were previously injected with Alexa555-conjugated CTB and Alexa647-conjugated CTB to allow visualization of ipsi- and contralateral projections in each visual target. Scalebar: 1mm. LC-MS/MS, liquid chromatography coupled to tandem mass spectrometry; SCN, suprachiasmatic nucleus; LGN, lateral geniculate nucleus; SCol, superior colliculus.

We based our analysis on four independent biological replicates of each RGC brain target. MS-based proteomic characterization revealed the expression of more than 3000 protein hits in each target with high confidence (ie detection of at least two specific peptides per hit, peptide-spectrum matching false discovery rate (FDR) < 1%). 3429 proteins were detected in the chiasm, 3128 in the SCN, 3660 in the vLGN, 3890 in the dLGN and 4140 in the SCol (**Table 1**). We verified the high reproducibility of our experimental design by plotting the extracted abundance of each protein hit across all replicates (**Supplementary Figure 1**). Focusing on the 3000 most abundant proteins from each brain region (as ranked by abundance), a gene ontology (GO) analysis conducted with DAVID revealed a high enrichment in proteins related to myelin sheath (GO:0043209), mitochondrial constituents (eg, GO:0005759 mitochondrial matrix, GO:0005743 mitochondrial inner membrane) and synapses (eg, GO:0008021 synaptic vesicle, GO:0014069 postsynaptic density), all terms characteristic of neuronal activity (**Figure 2, Supplementary Figure 2**). Interestingly, some of the most enriched GO terms shared by all regions of interest are linked to cell adhesion (eg, GO:0005913 cell-cell adherens junction, GO:0005925 focal adhesion, GO:0098641 cadherin binding involved in cell-cell adhesion, GO:0098609 cell-cell adhesion), reflecting the high connectivity between the neuronal and glial partners in the different brain targets, as well as tissue integrity (**Figure 2**). Identification of these enriched adhesion-related GO terms is based on the detection of adhesion proteins such as L1CAM, which have been shown to be critical for guidance of RGC axons during development of the visual system (Brittis and Silver, 1995) and of neonatal CST axons (Castellani et al., 2000).

**Figure 2:**
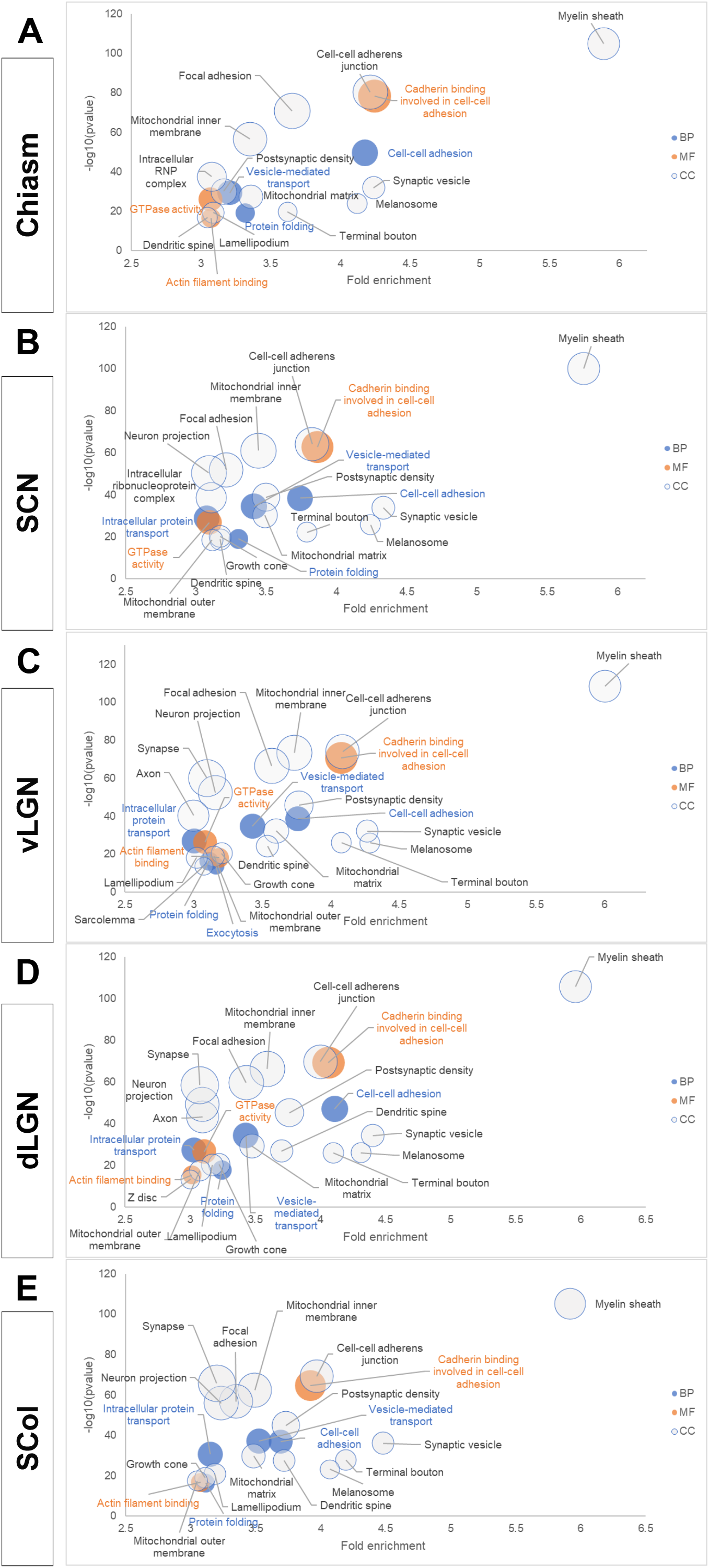
Adult visual targets exhibit enrichment of proteins related to connectivity and synaptic activity. **(A-E)** Bubble plots displaying enriched Gene Ontology (GO) terms in each visual system target, as analyed with DAVID. Terms associated with biological processes (BP) are represented in blue, cellular compartments (CC) in grey and molecular functions (MF) in orange. Only GO terms with a protein count higher than 50, a fold enrichment higher than 3 and a corrected p-value < 0.01 are represented. The bubble size of each GO term is representative of the number of proteins detected in the corresponding brain region.

### Adult visual targets express guidance and guidance-associated factors

We next aimed to characterize the molecular environment of adult neurons to understand whether this could affect their capabilities to be guided when regenerating. For that, we focused on proteins whose function or expression is related to adhesion, extracellular matrix, axon growth and guidance, or glia (**Table 2, Figure 3**). In all datasets, 121 proteins related to cell adhesion were identified, among which 70 proteins were present in all targets (58%) (**Table 3, Figure 3A**). Almost 80% proteins detected in the vLGN were detected in the dLGN, consistent with the fact that these two nuclei are anatomically close (**Supplementary Figure 3A-B**). These include well-known adhesion proteins, such as the Neural cell adhesion molecule Ncam1 (**Figure 3B**) known to induce neurite growth (Ditlevsen and Kolkova, 2010; Pollerberg et al., 2013) and Ncam2 recently shown to be required for growth cone formation (Sheng et al., 2015). Another example is APLP1, homologue to the amyloid precursor protein (APP) synaptic adhesion molecule, which was also detected in all selected targets, owing to its role in synapse maintenance and activity (Schilling et al., 2017). Members of the synaptic cell adhesion molecule family Cadm2, Cadm3 and Cadm4 were also identified in all targets. These cell adhesion molecules were demonstrated to be involved in axon pathfinding for midline crossing in the developing spinal cord (Frei et al., 2014). More specifically, the protein Neurofascin shown to stabilize synapses via transsynaptic contact stabilization (Kriebel, 2011) was detected in the LGN only. Together, these results highlight the adhesive nature of the adult intact brain probably reflective of the functional synaptic connectivity. Their expression in the intact visual targets provides information on the molecular environment that could challenge adult neuron regeneration.

**Figure 3:**
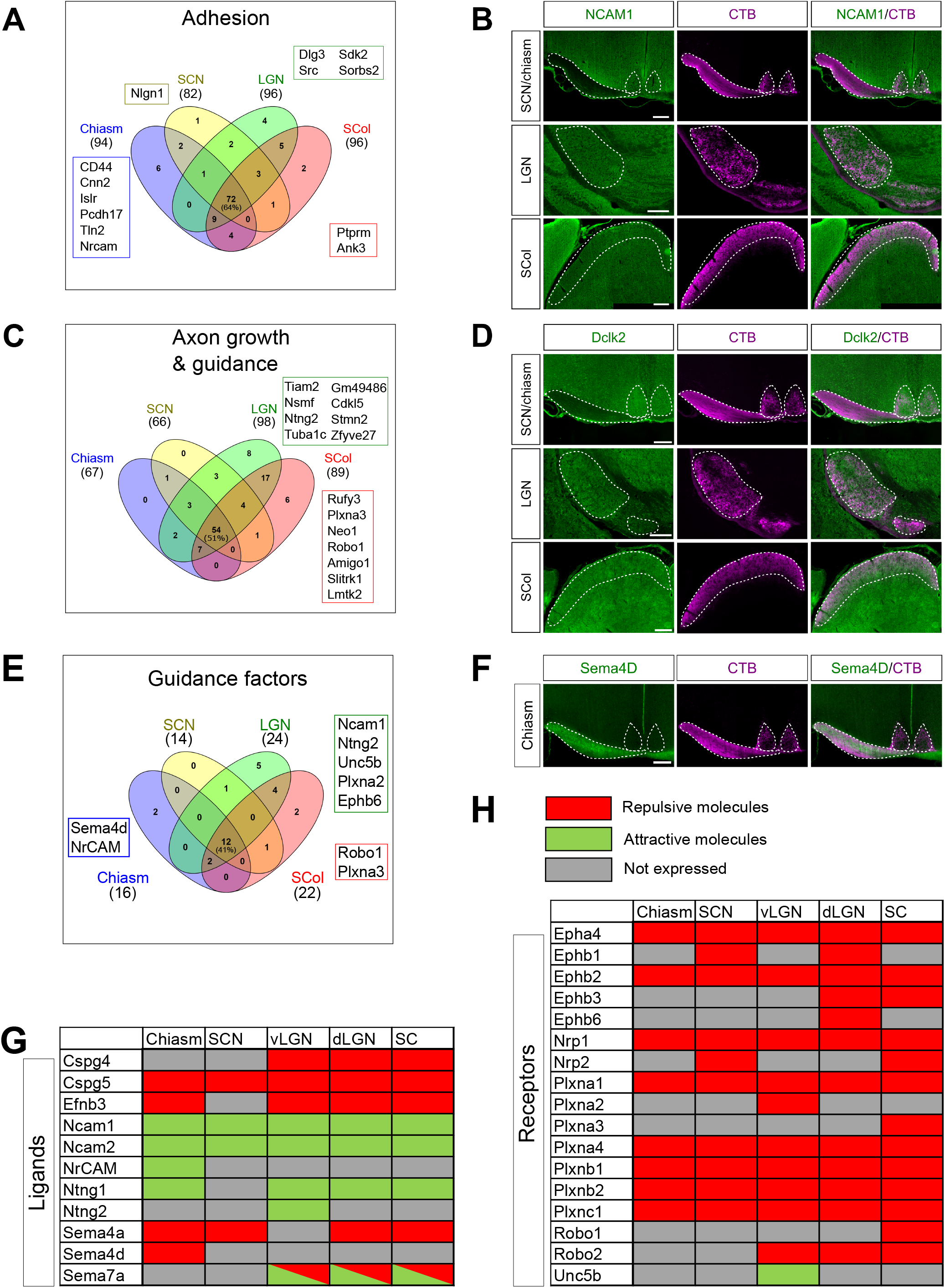
Adult visual targets express adhesion- and axon growth and guidance-related proteins. **(A)** Venn diagram representing the shared adhesion-related hits (number of proteins and percentage of the protein number in the adhesion category as defined in **Table 2**). In boxes are indicated proteins uniquelly detected in each visual target. **(B)** Immunofluorescent labelling of Ncam1 expressed in each visual target in an adult intact brain (colabelled with CTB). **(C)** Venn diagram representing the shared axon growth and axon guidance-related hits (number of proteins and percentage of the protein number in the axon growth and guidance category as defined in **Table 2**). In boxes are indicated proteins uniquelly detected in each visual target. **(D)** Immunofluorescent labelling of Dclk2 expressed in each visual target in an adult intact brain (co-labelled with CTB). **(E)** Venn diagram representing the shared guidance ligands and receptors (number of proteins and percentage of the protein number in the guidance ligands and receptors category). In boxes are indicated proteins uniquelly detected in each visual target. **(F)** Immunofluorescent labelling of Sema4D expressed in the optic chiasm (co-labelled with CTB). **(G)** Table representing the detection (green for attractive molecules, red for repulsive molecules) or the absence of detection (grey) of guidance ligands in each brain target. **(H)** Table representing the detection (green for attractive molecules, red for repulsive molecules) or the absence of detection (grey) of guidance receptors in each brain target. Scalebars: 200µm.

Another class of molecules of interest are extracellular matrix and ECM-related components. The ECM is an essential structural component of the adult brain and spinal cord. It plays a key role in the regulation of memory formation and plasticity, and acts as a repulsive structure of the glial scar following CNS injury (Benarroch, 2015; Fawcett, 2015; Silver and Miller, 2004). Components of the ECM such as chondroitin sulfate proteoglycans (CSPG) are known to play a growth-inhibitory role on injured axons, thereby hindering regeneration potential following CNS injury (Silver and Miller, 2004). Recently, the ECM has also been described as a regulator of guidance events during development (Donato et al., 2018; Schaeffer et al., 2018; Wiese and Faissner, 2015). The ECM can play a role of molecular scaffold of guidance molecules. Interestingly, molecules of the ECM can also modulate the guidance activity of such molecules, for example by enhancing the repulsive activity of Sema3A (Dick et al., 2013; Schaeffer et al., 2018). In our study, 57 proteins of the ECM were identified (**Table 3, Supplementary Figure 3C**). Among them, 24 (42%) were shared between all the targets, including members of the collagen (Collagen alpha-1 (IV) chain, collagen alpha-2 (IV) chain, collagen alpha-1 (XII) chain), the laminin (Laminin subunit alpha-5, laminin subunit beta-2) and the proteoglycan (hyaluronan and proteoglycan link protein 1, 4 Hpln1 and Hpln4, chondroitin sulfate proteoglycan 4 and 4, Cspg4 and Cspg5) families. We found Cspg4 expression in the LGN and SCol (**Figure 3G, Supplementary Figure 3C-D**). Cspg4/NG2 inhibits axon growth in vitro (Lee et al., 2013), but although it is yet unclear whether NG2-expressing cells, a major cellular component of the glial scar, provide in fact a stabilized substrate to support axon growth after spinal cord injury (Filous et al., 2014; Sellers et al., 2009). Another prominent hit is the ECM glycoprotein Tenascin-C detected in all targets. Tenascin-C displays bidirectional activity during development, depending on neuronal subtype and integrin receptor expression (Husmann et al., 1995). Its upregulation in the glial scar following CNS injury seems to be associated with a growth-promoting, integrin-dependent activity (Cheah et al., 2016; Chen et al., 2010). More specifically, the SCN displays detection of heparan sulphate proteoglycan 2 (Hspg2), which plays a role in growth, migration and repulsive axon guidance (Yamaguchi, 2001). Here, our datasets highlight the rich ECM environment of the target regions of RGC axons, which is of particular relevance to a context of axon growth and guidance.

Next, we looked at proteins associated with glial cells, known to impact axon growth during development (Silver et al., 1993) and in adult following injury (Fitch and Silver, 1997; Silver and Miller, 2004). These include myelin components and oligodendrocyte-associated proteins, for example the potent neurite inhibitory molecule Nogo-A (Reticulon-4A) (Chen et al., 2000; GrandPré et al., 2000); astrocyte markers such as GFAP (**Table 3, Supplementary Figure 3E-F**) and GLAST; microglia markers such as CD11b (also known as Integrin alpha-M). Glia-associated proteins showed little region specificity (71% of glia-associated molecules detected in all targets). This result is reflective of an environment highly enriched in bundles of myelinated axons (**Supplementary Figure 3E**).

Furthermore, we analysed proteins related to axonogenesis, axon extension, cytoskeleton, growth cone and axon guidance, a subset of proteins that we termed axon growth and guidance. We found expression of 109 proteins, including 57 (47%) shared by all visual targets (**Table 3, Figure 3C**). The protein doublecortin-like kinase 2 (Dclk2), recently shown to promote axon growth via induction of growth cone reformation (Nawabi et al., 2015), is expressed in all visual targets (**Figure 3D**). Axon growth and guidance proteomes of the chiasm and the SCN are very similar, which could be explained by their close anatomical proximity. On the other hand, the SCol and the LGN display unique detection of proteins related to axon growth and guidance and share expression of 19 growth and guidance molecules (**Table 3, Figure 3C**). Among them, several guidance factors were detected such as Robo2, EphB3, RGMA (Repulsive guidance molecule A), and Slik5 (Slit and NTRK-like family member 5). Although originally described as regulators of axon navigation in the developing nervous system, these guidance factors persist throughout adult life and are implicated in physiological processes such as synapse maintenance and strengthening (Sun et al., 2018), in particular at the closure of the critical plastic period of development (Fawcett, 2009). On the other hand, in the context of regeneration, the presence of repulsive cues may impair the growth capacity of regenerating axons and/or misguide them away from their proper targets. For example, RGMa is also expressed in the SCol and LGN and has been described to mediate glial scar formation after a stroke (Zhang et al., 2018), indicating that it could constitute a physical barrier to axon regeneration.

### Classical guidance cues and receptors are expressed in the intact brain

We focused further our research on classical guidance ligands and receptors of the Semaphorin, Slit, Ephrin and cell adhesion molecule (CAM) families (**Figure 3E**). These molecules are originally described for their guiding role of navigating axons during development. Several guidance molecules have also been described in the adult to play a role in physiological conditions such as during synapse functioning (Bouzioukh et al., 2006, 2007; Glasgow et al.; Kania and Klein, 2016). Very interestingly, our datasets revealed the presence of classical guidance cues and associated receptors in visual targets of the intact adult brain. Indeed, we found that 10 guidance ligands and 17 guidance receptors (thereafter designated as “guidance factors”) are expressed in the visual system primary targets in adult (**Figure 3G-H**). 12 guidance factors were detected in all targets. Among those common proteins, some are permissive to axon growth and guidance such as the immunoglobulin cell adhesion molecules Ncam1 and Ncam2 (Walsh and Doherty, 1997).

However, others prevent the axon navigation such as Cspg5 which is a proteoglycan of the ECM that play a role of barrier molecule affecting axon growth (Laabs et al., 2005). Sema4A is also present in all the brain targets. It induces cell morphological changes notably growth cone collapse by interacting with PlexinB receptors (Yukawa et al., 2005, 2010). Interestingly, PlexinB1 and PlexinB2 are receptors present in all the targets, suggesting a physiological role for the interaction Sema4A/PlexinB in the brain. Another notable example is the expression of Eph receptors such as EphB2 and Ephrin ligands such as EphrinB3 (Efnb3). EphB2 has been shown to modulate excitatory synapse density and activity via trans-synaptic interaction with EphrinB3 (Kayser et al., 2006; McClelland et al., 2010). Notably, Efnb3 has been described for its role as a repulsive midline barrier for many ipsilateral axons in the developing spinal cord (Chédotal, 2019; Imondi et al., 2000).

Interestingly, some guidance ligands show unique detection in specific targets. This is the case for Sema4D in the chiasm (**Figure 3F**). Sema4D is known to induce cell morphological changes notably growth cone collapse by interacting with PlexinB1 (Swiercz et al., 2002; Tasaka et al., 2012). Sema4D has been recently shown to be upregulated in ipsilateral RGC during optic chiasm decussation (Wang et al., 2016). This suggests that if regenerating axons express PlexinB1, they might be repelled by Sema4D expression in the chiasm, therefore causing misguidance and incorrect ipsi-vs contralateral projection. This phenotype has been observed in different models of long-distance regeneration in the optic nerve (Belin et al., 2015; Lim et al., 2016).

### Optic nerve injury causes some modifications in the proteome of the visual targets

In order to understand the influence of the injury on visual targets proteomes, we performed bilateral optic nerve crush in adult (6 week-old) mice (**Figure 4A**). Bilateral optic nerve crush is required to obtain a full denervation of all visual targets, as each target is innervated by both eyes (about 5% RGC projecting ipsilaterally, about 95% contralaterally in mouse). We collected the brain targets four weeks later, long term after the injury and at a time point where regenerative axons reach the visual target regions in long-distance regeneration models (Belin et al., 2015; Lim et al., 2016). Samples from visual targets at 28 days post-crush (28dpc) were compared with the intact condition by MS-based label-free quantitative proteomics (**Figure 1A**), using four biological replicates for each condition. Efficiency of optic nerve crush was verified by CTB injection two days prior tissue dissection (**Figure 4B**). Biological replicates of the crush condition showed high consistency as shown by the scatterplots of protein abundances (**Supplementary Figure 1**). For each brain target, principal component analysis (PCA) showed clustering of replicates according to the condition (**Supplementary Figure 4A-E**). Very interestingly, the injury leads to modification of the proteome of the visual targets, with 311 proteins differentially expressed in the optic chiasm, 45 in the SCN, 26 in the vLGN, 92 in the dLGN and 92 in the Scol (log_2_ fold change > 0.8, FDR < 5%) (**Table 4, Figure 4C-D, Supplementary Figure 4F-H**). We were surprised to unravel so many differences in protein expression in targets anatomically far from the injury site, in the chiasm (about 5mm from the eye ball in mouse) and also in more distal regions such as the dLGN and the SCol.

**Figure 4:**
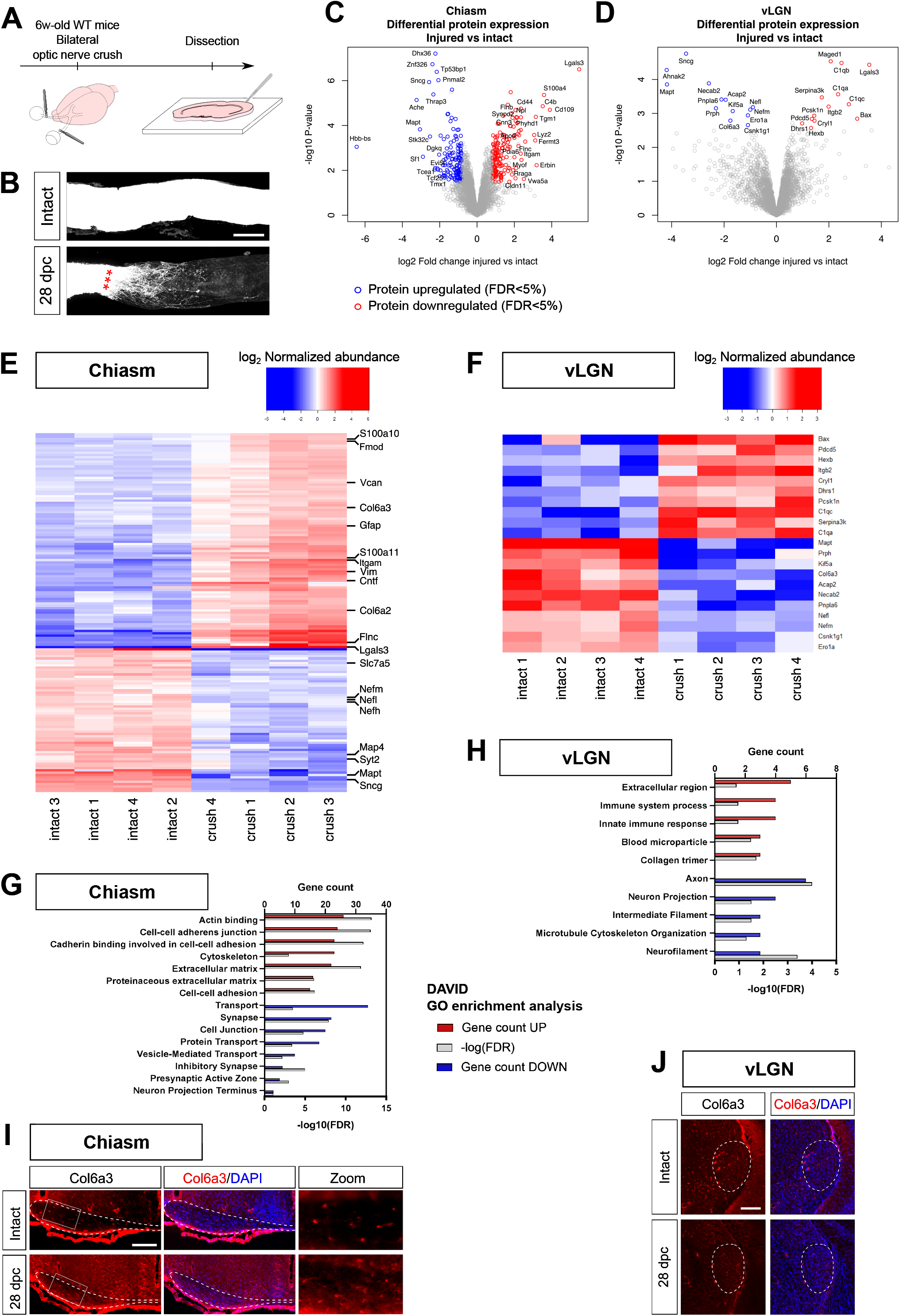
Optic nerve crush causes modifications in the proteome of adult visual targets. **(A)** Experimental design and timeline. 6 week-old mice undergo bilateral optic nerve crush and brains were dissected 28 days after the injury (28 dpc). **(B)** Transparized optic nerve with CTB labelling in intact and injured conditions. **(C-D)** Volcano plots showing differentially expressed proteins in the optic chiasm and in the vLGN, in injured versus intact conditions. Highlighted are protein hits with log_2_ fold change < 0.8 and FDR < 5%. **(E-F)** Heatmaps showing differentially expressed proteins between injured (crush) and intact conditions. The values are abundance normalized across all samples for each protein hit. For the optic chiasm, proteins with FDR < 1% are represented. For the vLGN, proteins with FDR < 5% are represented. **(G-H)** GO terms associated with lesion-modulated hits in the optic chiasm and in the vLGN. In red is represented the protein count of the corresponding GO for up-regulated proteins. In blue is represented the protein count of the corresponding GO for down-regulated proteins. In grey is represented the significance as -log_10_(FDR). **(I-J)** Immunofluorescence showing the expression of Col6a3 in the optic chiasm and in the vLGN, in intact and injured conditions.

Our datasets reveal a high number of differentially expressed proteins in the chiasm (311), among which 165 are upregulated and 146 are downregulated after the injury (**Table 4, Figure 4C**). We submitted the list of differentially expressed proteins to a functional classification analysis with DAVID. Interestingly, proteins upregulated after injury are associated with cell adhesion and extracellular matrix (eg Talin-1 (TLN1) and Galectin-3 (LEG3)) (**Figure 4E-G**), supporting the hypothesis of a remodelling of the environment of the intermediate target after injury. On the other hand, proteins downregulated, such as Synaptotagmin-2 (SYT2) and Exportin-T (XPOT), are associated with synapse and transport (**Figure 4E-G**), consistent with the alteration of the physiological features of intact axons 28 days post-crush. For example, we found an upregulation of Collagen VI alpha 3 (Col6a3) in the chiasm following injury (**Figure 4I**).

We found a smaller number of differentially expressed proteins in the vLGN (26), with 13 going up and 13 going down after the injury (**Figure 4D**). DAVID analysis allowed us to highlight a robust enrichment of terms related to inflammatory response and to collagen in the list of proteins upregulated. Conversely, we found a downregulation of structural components of the axon, for example the microtubule-associated protein tau (Mapt) involved in the stability of microtubules in the axon (**Figure 4F-H**). These surprising results suggest that after axonal injury, denervated targets may respond to the injury even at a distance from the lesion. The local changes in inflammatory proteins (Complement C1q subcomponents (C1QA, C1QB, C1QC) upregulation) observed in distal targets may be a consequence of RGC axon degeneration (glial activation, debris clearing) or the protein remodelling of the denervated targets themselves in response to the injury. Interestingly, the ECM component Col6a3 that is upregulated in the chiasm was found downregulated in the vLGN after injury (**Figure 4J**). Altogether, these modifications reveal a dynamic environment in the adult brain, with some specificity among the different brain targets. These changes influence the capacity of regenerative axons to resume developmental processes and reconnect their proper target.

### Guidance molecules in the brain are not modified following injury

In most models of regeneration, only few axons are observed to enter the brain target (Lim et al., 2016). In fact, most regenerating axons seem to avoid the brain nuclei. Interestingly, the presence of repulsive guidance molecules in the adult visual system even after the injury could explain the impossibility for regenerating axons to reconnect their proper target. Our data show that the injury modulates the expression of many proteins even in distal targets of the visual system. Regarding canonical guidance factors, we observed that their level of protein expression rmains steady in the adult visual targets. Importantly, guidance factors which were described mostly during development, are still expressed in the adult optic chiasm and RGC targets and could potentially still have guidance activity.

We verified these results by performing Western Blot analysis on independent biological replicates of each brain target in intact versus injured condition (**Figure 5**). Focusing on several guidance factors and regulators, such as ECM components, cell adhesion molecules and guidance fators, we confirmed expression in intact condition and found no difference of the level of expression between intact and injured. The extracellular matrix molecule TNC is expressed in the optic chiasm with no significant variation between intact and injured conditions (**Figure 5A**). The cell adhesion molecule Ncam1 known to have a potent axon growth-promoting activity is expressed in all brain targets (SCN, LGN, SCol) and its expression is not modified after injury (**Figure 5B-E**). Similary, the guidance cue Sema7a shows a steady expression in the LGN and SCol (**Figure 5C-E**). These results indicate that the lesion does not modify the guidance landscape of the brain target. Guidance factors stably expressed in the adult brain may be the cause of guidance defects observed in regenerative models after injury.

**Figure 5:**
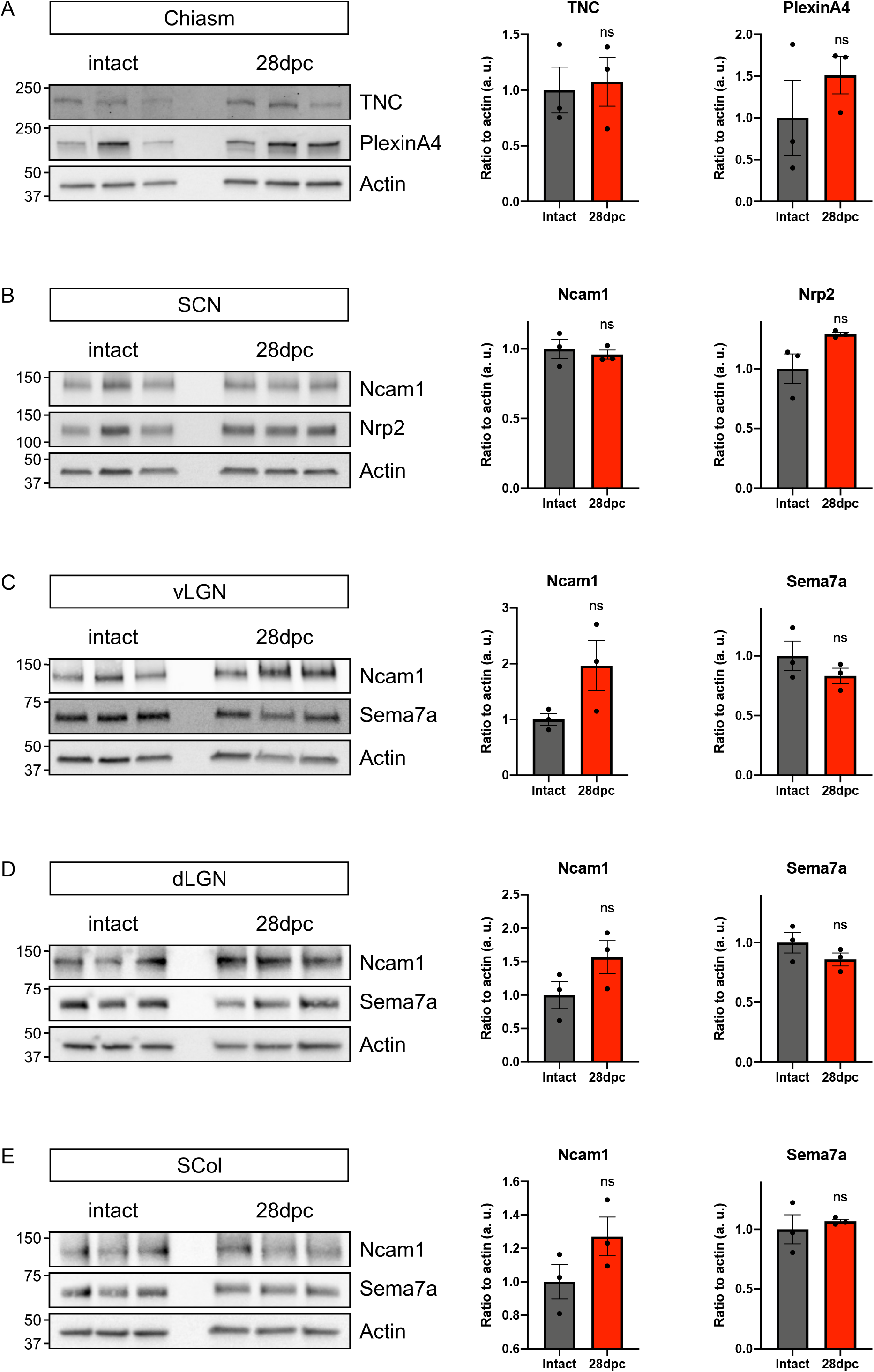
Guidance and guidance-associated factors remain steady in the adult visual targets after optic nerve injury. **(A-E)** Western blot analysis of selected proteins in each visual targets (left) and corresponding quantification (right). Each lane corresponds to tissue collected from one animal. For each sample, protein expression is quantified by pixel densitometry relative to actin and normalized to intact condition. Student’s t-test, ns: not significant.

As many guidance molecules were detected in the brain targets, we asked then whether RGC can integrate these signals. Therefore, we looked at the expression of corresponding guidance receptors in RGC. As proteomic data are more difficult to obtain due to limited amound of material in FACS-sorted RGC, we used available transcriptomic datasets of two recent single-cell studies (Rheaume et al., 2018; Tran et al., 2019) and of two regenerative models: co-deletion of Pten and SOCS3 (Sun et al., 2011) and overexpression of Sox11 (Norsworthy et al., 2017). Using our map of guidance ligands expressed in mature visual targets (**Figure 3G**), we explored expression of the corresponding guidance receptors. We found that corresponding receptors are expressed in RGC both in intact WT condition and in regenerative (post-injury) condition. For example the semaphorin 4 receptor Plexin-B2 is expressed is RGC in intact and in regenerative (post-crush) conditions (**Supplementary Figure 5A**). On the other hand, the transcript for the semaphorin 4 receptor Plexin-B3 is not detected in either intact datasets, and detected only in Pten-/- SOCS3-/- regenerative RGC (Sun et al., 2011). In the intact datasets, we highlighted the number of RGC clusters showing expression of the receptors of interest. Interestingly, despite compelling evidence of differential injury response of neuronal subpopulations (Duan et al., 2015; Tran et al., 2019), almost all RGC clusters express guidance receptors for CSPG family members, Ephrin-B3, Ncam1 and 2, Sema4d and Sema7a, suggesting that adult guidance response is shared by all RGC subpopulations. Furthermore, we looked at the variation of axon guidance molecules between regenerative conditions and non-regenerative (WT) conditions following injury (Norsworthy et al., 2017; Sun et al., 2011). Very interestingly, we found that many guidance receptors and cell adhesion molecules are dynamically regulated (either up or down) in the regenerative conditions, eg Robo1 and Plexin-A1 strongly upregulated compared to WT, which suggests that regenerative RGC modify their guidance response potential in these models (**Supplementary Figure 5B**). We also observed that some guidance receptors, such as EphA7 and EphA3, vary in the opposite directions in the two regeneration models. This is most probably due to the differences in pathway activation and accounts for the complexity of RGC guidance response that has to be considered to correct their misguidance.

### The guidance landscape is established during innervation of brain targets

Our datasets show that in adult the injury and denervation of visual targets does not modify the expression of guidance factors. We wondered whether establishment of this guidance map is concomitant with the initial innervation of the target regions during development. We focused on the LGN as a proof-of-concept and looked at the different timepoints of its innervation by retina ganglion cells. We looked at the chondroitin sulfate proteoglycan CSPG4/NG2, inhibitory for axon growth in vitro (Lee et al., 2013), that we found expressed in the adult LGN (**Supplementary Figure 3D**). To track innervation of the LGN during development, we injected one eye of P0 to P14 mouse pups were injected with CTB555. Contralateral RGC axons enter the LGN between E16 and P0, while ipsilateral RGC axons enter between P0 and P2, with synapse refinement and eye-specific segregation of innervation territories happening at eye opening (around P12) (Guido, 2018). Strikingly, the expression of CSPG4 is very low in the LGN until P4, and starts to increase from P6 on up to P14 (**Figure 6**).

**Figure 6:**
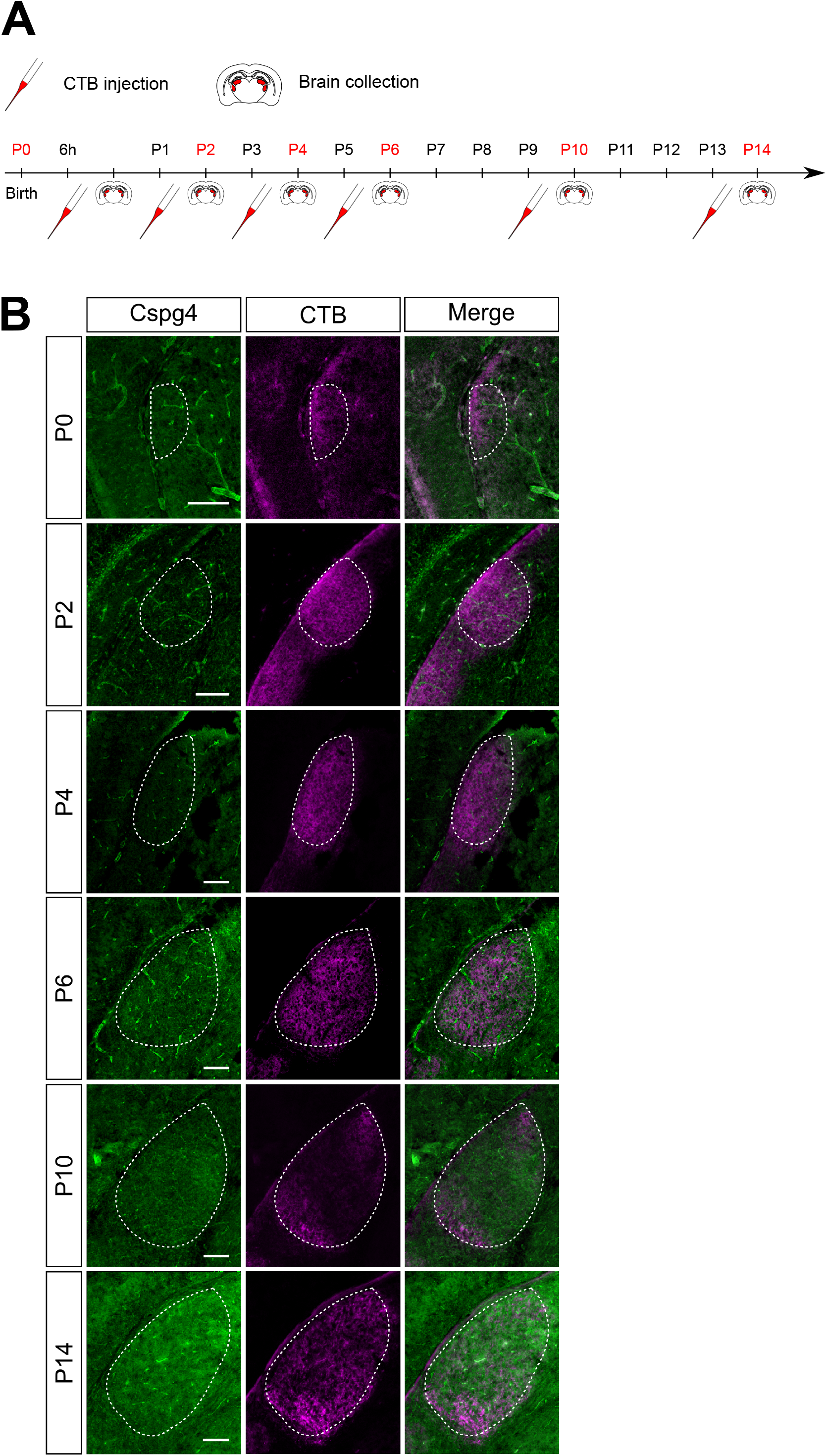
The guidance map is established during innervation of the developing visual target regions. **(A)** Experimental design and timeline. For P2 to P14 timepoints: one day before sample collection, one eye of mouse pups were injected intravitreally with CTB-555. For P0 timepoint: 6h before sample collection, one eye of mouse pups were injected intravitreally with CTB-555. (B) Immunofluorescence showing expression of CSPG4 in the developing dLGN (co-labelled with CTB). Scalebar: 100µm.

These observations show that at the time of innervation, the guidance cue CSPG4 is absent, while axons are able to enter the LGN. Then its expression increases when synapses are formed and need to stabilize. CSPG upregulation correlates with the stabilization of the circuit and its maintenance. The reverse timing of expression is observed with the CSPG Aggrecan expressed in the LGN at P0 and progressively degraded by proteases to allow innervation of corticogeniculate axons at P7 (Brooks et al., 2013). More generally, CSPG are a major component of the perineuronal nets and their upregulation during development coincides with the closure of the critical plastic period during circuit consolidation (Carulli and Verhaagen, 2021). Here, we find that CSPG4 is not only maintained through adulthood, it is also present following injury. This way, it may prevent reinnervation of regenerative RGC axons through a possibly repulsive activity.

Altogether, our results suggest that there is the tight spatio-temporal regulation of guidance cues expression the coincides with a window allowing the innervation of the brain targets. During adulthood, guidance cues may switch their function over time, from a canonical axon pathfinding activity to synaptogenesis during circuit formation, then to a structural and regulatory function of synaptic activity in adult (**Figure 7A**). Following injury, these factors may resume their guidance activity and contribute to misguidance of regenerative axons and/or prevent reinnervation of the correct targets, leading to absence of functional reconnection. Our proteomics data allow an extensive characterization of adult visual targets, which is indispensable to understand the guidance landscape of regenerative axons (**Figure 7B**).

**Figure 7:**
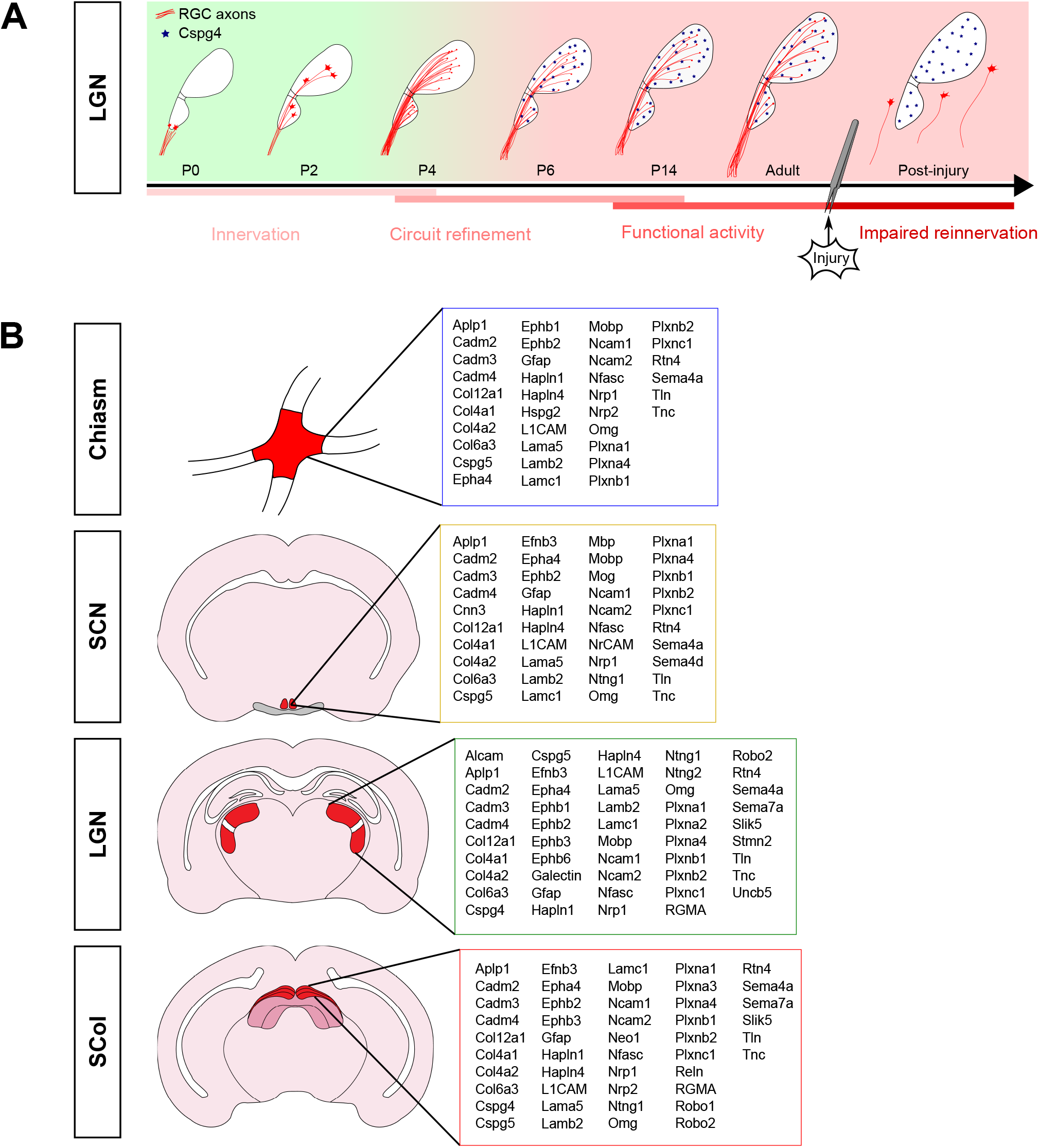
Guidance maps in the adult brain targets are set during development and provide insight into the guidance defects encountered by regenerative axons. **(A)** Using the expression of CSPG4 in the dLGN as a proof-of-concept, our data suggest that the spatio-temporal regulation of guidance cues contribute to establishment of the neuronal circuit during development. In adult, guidance cue expression remain and display physiological functions. Following optic nerve injury, visual brain targets get denervated and undergo proteome remodelling. Guidance cue expression remain steady and may contribute to misguidance of regenerative RGC away from their initial target. (B) Summary of guidance maps as described in the present proteomics study for each brain target of the adult visual system. SCN, suprachiasmatic nucleus; LGN, lateral geniculate nucleus; SCol, superior colliculus.

## DISCUSSION

In this study, we performed an extensive proteomic description of the primary targets of the visual system innervated by RGC axons in the intact adult brain and after bilateral injury of the optic nerves. Axons of the adult CNS fail to regrow spontaneously because of the lack of intrinsic regenerative molecular programme (He and Jin, 2016). However, even long-distance regeneration models show poor functional recovery, particularly because regenerative axons display guidance defects that prevent reconnection to their proper target (Luo et al., 2013; Belin et al., 2015; Pernet et al., 2013a, 2013b). In this context, we sought to determine the protein content of the RGC primary targets to understand if the molecular environment could contribute to this misguidance. Despite recent advances in transcriptomic profiling of neuronal populations down to the single-cell level (Kalish et al., 2018; Rheaume et al., 2018; Tran et al., 2019; Wen et al., 2020), extensive characterization of the proteome in the adult CNS is still needed. For this reason, we explored the proteomes of RGC targets: the optic chiasm, a critical intermediate target, and the major functional nuclei of the visual system: the SCN, the ventral and dorsal LGN and the superior colliculus.

Previous proteomics analysis on the supra-chiasmatic nucleus has been conducted with a particular focus on the cyclic expression of proteins that is directly correlated to the regulation of the circadian clock (Chiang et al., 2014; Deery et al., 2009). Here, we focused on the connectivity of the circuit with a specific interest in guidance and guidance-associated factors. We found several proteins expressed by all targets reflective of the shared guidance signature of RGC targets (eg Cspg5, Ncam1 and 2, members of the Plexin family). We also uncovered guidance factors that were uniquely detected in specific targets, eg Sema4D in the optic chiasm. In the mature CNS, canonical guidance cues are involved in physiological processes, such as plasticity and regulation of synaptic activity, for example Eph/Ephrin signaling (Kania and Klein, 2016). As the system matures, these guidance factors may act as a roadblock to progressively limit neuronal plasticity and strenghten the neuronal circuit. Importantly, this brings up the possibility of an active influence of such cues on guidance of regenerative axons and impairment of functional reinnervation at the adult age. Failure of CNS regeneration has long been attributed to the expression of growth-inhibitory substances at the lesion site. Our datasets provide a guidance map in the adult CNS where guidance and guidance-associated factors do not simply block axon regrowth, but instead guide them away from the correct track.

To explore this hypothesis further, we looked at proteome changes in the RGC targets upon denervation. Our datasets reveal that bilateral optic nerve crush results in modification of protein expression. Recently, protein expression changes after axon injury have been explored in the RGC soma (Belin et al., 2015), revealing key signalling hubs of the neuronal injury response. In our study, we were interested inn the neuronal injury response of the primary targets of the lesioned RGC. Intriguingly, we discovered protein expression changes after injury, even in distal targets. Multiple, non-exclusive hypotheses arise here: i) target neurons are not sustained anymore by RGC neuronal activity undergo protein expression changes; ii) molecular changes of the distal response may be intrinsic to these target neurons and possibly consequent to forward signalling of axon injury; iii) axon degeneration down to the lesion site triggers a local inflammatory response which in turns causes injury response of the neuronal and non-neuronal populations with adaptation of protein expression. It is more expected that the latter is a transient response short to mid-term following the injury, as observed in the Wallerian degeneration model (Llobet Rosell and Neukomm, 2019). In our case, at 28dpc, we expect that axon death and debris clearance by glia has occurred. Yet, sustained activation of the glia and activation of the immune system are a feature of CNS injury (Allison and Ditor, 2015; Burda and Sofroniew, 2014; Jassam et al., 2017), consistent with what we found for example in the vLGN with enrichment of immune response in upregulated proteins.

What do we know about molecular modifications in neuronal targets of lesioned axons? Previous transcriptomic work focused on gene expression changes in the superior colliculus in the course of development and following injury (Liu et al., 2010a; Mei et al., 2013). This study highlighted functional groups of genes regulated by the injury, including transport and metabolism. However, this work was based on monocular enucleation (Liu et al., 2010b), which is less specific than RGC axon injury. More recently, a transcriptomic profiling of the spinal cord target tissue after stroke allowed to highlight two phases of the target injury response: a primary phase characterized by the inflammatory response in the target area, and a secondary later phase characterized by secretion of connectivity-promoting growth factors (Kaiser et al., 2019). Such changes in the long-term might affect axon growth and guidance of regenerative axons and open a therapeutic window of modulation of gene expression to promote growth and synaptogenesis. Importantly, our datasets highlight a remodelling of the proteome of visual brain targets. On top of the challenge of visual target reinnervation by regenerative RGC, one should consider that these changes in protein expression may affect the functionality of the connectivity. In other words, if regenerative axons manage to enter the proper target and form synapses, will injury-induced modifications of the target itself impair its capacity to respond to neural activity and to resume functional connectivity? Is the plastic period definitely over in the adult lesioned CNS?

With the hope to achieve functional reconnection of regenerative fibers, it is essential to characterize the cellular environment to tackle guidance defects, in particular in the optic chiasm where the majority of regenerative fibers get lost before entering the optic tracts (Belin et al., 2015; Crair and Mason, 2016; Luo et al., 2013; Pernet et al., 2013b, 2013a). The strength of our approach is to give an unbiased characterization of the protein content of the optic chiasm and primary targets of RGC axons. For particular targets such as the optic chiasm, the pool of upregulated proteins relates to adhesion and extracellular matrix, suggestive of a remodelling of the environment that might affect guidance in the case of regeneration. In addition, our results show that i) numerous guidance factors are present in the intact mature brain and ii) these factors remain steady upon injury, which means their expression and possibly guidance function are maintained in the lesioned CNS.

Finally, it is important to consider the response of RGC themselves. In several long-distance regeneration models, genetic manipulation of molecular pathways in neurons may affect the expression of proteins of the growth and guidance machinery. It cannot be excluded that rather than being actively misguided by guidance factors expressed in targets of interest, regenerative RGC axons are in fact unresponsive to pathfinding. Altogether, our datasets provide an unbiased extensive protein screen of primary visual targets of the adult CNS, in intact and in injured conditions. Many guidance and guidance-associated factors could be identified, that may influence the target-tracking activity of adult regenerating axons. Their physiological function and their implication in preventing connectivity remain to be determined. In this context, manipulation of expression of the corresponding receptors in RGC opens up great promises in the reinnervation of the functional targets in the lesioned adult CNS.

## MATERIAL AND METHODS

### Mice

Wild-type (WT) pups (P2, P4, P6, P10, P14) and adult (6 week-to 10 week-old) mice were used in this study, regardless of sex. All the in vivo experiments were performed in accordance with our ethics protocol approved by the institution, local ethics committee and the French and European guidelines (APAFIS#9145-201612161701775v3 and APAFIS#26565-2020061613307385v3).

### Optic nerve crush injury

Optic nerve crush was performed as described before (Schaeffer et al., 2020). Briefly, 6 week-old mice were anesthetized with intraperitoneal injection of ketamine (60–100 mg/kg) and xylazine (5–10 mg/kg). A mini bulldog serrefines clamp was placed to display the conjunctiva. The conjunctiva was incised lateral to the cornea. The refractor bulbi muscles were gently separated and the optic nerve was pinched with forceps (Drumont #5 FST) at 1mm from the eyeball during 5 seconds (Park et al., 2008; Schaeffer et al., 2020).

### CTB injection

One day before sample collection, mice were anesthetized as described before and intravitreal injections of CTB-555 (Cholera toxin subunit B, Alexa Fluor 555-conjugated, ThermoFisher Scientific) were performed. The external edge of the eye was clamped using a mini bulldog serrefines clamp (FST) to display the conjunctiva. 1µl of CTB-555 (1mg/mL) was injected into the vitreous body using a glass micropipette connected to a Hamilton syringe (Park et al., 2008; Schaeffer et al., 2020).

### Sample collection

Mice were subdivided into 2 groups: control (intact mice and mice that underwent optic nerve injury. 10 weeks old mice from both groups were anesthetised using isoflurane (Equipement Vétérinaire Minerve). After cervical dislocation, eyeballs and brains were dissected out. Using a vibratome (Leica VT1200S) 300µm fresh brain sections were collected in ice-cold Hibernate A (ThermoFisher Scientific). Chiasm, suprachiasmatic nucleus (SCN), ventral lateral geniculate nucleus (vLGN), dorsal lateral geniculate nucleus (dLGN) and superior colliculus (SCol) were micro-dissected under a binocular microscope (Zeiss SteREO Discovery.V8) and flash frozen in dry ice. Eyes of control mice were previously injected intravitreally with CTB-555 to facilitate detection of the visual targets.

### Sample preparation

Total protein lysates were obtained by extraction in Laemmli 2x (recipe) and 15min incubation on ice. 0.5µl of benzonase (Sigma-Aldrich) was added to each sample and incubated for 10 min at 37°C to digest DNA. After denaturation at 95°C for 5 min, the protein concentration was determined using Pierce BCA Protein Assay Kit (ThermoFisher Scientific). To ensure sample quality and consistent quantification, a silver staining was performed by loading 1µg of proteins onto a 4-15% precast gel (Biorad). After electrophoresis, silver staining of the proteins was performed using the Silver Quest Kit (ThermoFisher Scientific).

### Mass spectrometry-based proteomic analyses

Proteins from tissue preparations were solubilized in Laemmli buffer before being stacked in the top of a 4–12% NuPAGE gel (Life Technologies), stained with R-250 Coomassie blue (Bio-Rad) and in-gel digested using modified trypsin (sequencing grade, Promega) as previously described (Salvetti et al., 2016). The dried extracted peptides were resuspended in 5% acetonitrile and 0.1% trifluoroacetic acid and analyzed by online nanoliquid chromatography coupled to tandem mass spectrometry (LC–MS/MS) (Ultimate 3000 RSLCnano and the Q-Exactive HF, ThermoFisher Scientific).

#### Chiasma and SCN samples

Peptides were sampled on a 300µm 5mm PepMap C18 precolumn (ThermoFisher Scientific) and separated on a 75µm 250mm C18 column (Reprosil-Pur 120 C18-AQ, 1.9 µm, Dr. Maisch HPLC GmbH). The nano-LC method consisted of a 240min multi-linear gradient at a flow rate of 300nl/min, ranging from 5 to 33% acetonitrile in 0.1% formic acid.

#### dlGN, vLGN and SCol samples

Peptides were sampled on a 300µm 5mm PepMap C18 precolumn (ThermoFisher Scientific) and separated on a 200cm µPACTM column (PharmaFluidics, Ghent, Belgium). The nano-LC method consisted of a 360min multi-linear gradient at a flow rate of 300nl/min, ranging from 5 to 33% acetonitrile in 0.1% formic acid.

For all tissues, the spray voltage was set at 2 kV and the heated capillary was adjusted to 270°C. Survey full-scan MS spectra (m/z = 400–1600) were acquired with a resolution of 60 000 after the accumulation of 3 × 10^6^ ions (maximum filling time 200ms). The 20 most intense ions were fragmented by higher-energy collisional dissociation after the accumulation of 10^5^ ions (maximum filling time: 50ms). MS and MS/MS data were acquired using the software Xcalibur (Thermo Scientific).

### Mass spectrometry-based proteomic data processing

Data were processed automatically using Mascot Distiller software (version 2.7.1.0, Matrix Science). Peptides and proteins were identified using Mascot (version 2.6) through concomitant searches against Uniprot (Mus Musculus taxonomy, June 2020 version), classical contaminants database (homemade) and their corresponding reversed databases. Trypsin/P was chosen as the enzyme and two missed cleavages were allowed. Precursor and fragment mass error tolerances were set, respectively, to 10ppm and 25mmu. Peptide modifications allowed during the search were: carbamidomethylation (fixed), acetyl (protein N-terminal, variable) and oxidation (variable). The Proline software (version 2.0) (Bouyssié et al., 2020) was used to merge and filter results for each tissue separately: conservation of rank 1 peptide-spectrum match (PSM) with a minimal length of 7 and a minimal score of 25. Peptide-spectrum matching (PSM) score filtering is then optimized to reach a False Discovery Rate (FDR) of PSM identification below 1% by employing the target decoy approach. A minimum of one specific peptide per identified protein group was set. For each tissue separately, Proline was then used to perform MS1-based label free quantification of the peptides and protein groups from the different samples with cross-assignment activated. After peptide abundances normalization, protein abundances were computed as a sum of specific peptides abundances.

### Statistical analysis of mass spectrometry-based proteomic data

Statistical analysis was performed using ProStaR (Wieczorek et al., 2017) to determine differentially abundant proteins between intact and crushed conditions. Protein sets were filtered out if they were not identified in at least two replicates of one condition. Protein sets were then filtered out if they were not quantified across all replicates in at least one condition. Reverse protein sets and contaminants were also filtered out. After log2 transformation, POV missing values were imputed with slsa method and MEC ones with 2.5-percentile value of each sample. Statistical testing was conducted using limma test. Differentially-expressed proteins were sorted out using a log_2_(fold change) cut-off of 0.8 and a p-value cut-off allowing to reach a FDR inferior to 5% according to the Benjamini-Hochberg procedure.

### Data analysis

#### Scatterplots

Scatterplots of protein hits were obtained by plotting the protein abundances across replicates of a same visual target region in either intact or injured conditions.

#### Gene Ontology analysis

Gene Ontology (GO) analysis was performed using DAVID (Database for Annotation, Visualization and Integrated Discovery) Bioinformatics Resources (version 6.8). Bubble plots were obtained by submitting the list of the 3000 more abundant proteins (ranked by protein abundance, see **Table 1**) of each brain target to DAVID. GO terms were divided into 3 groups: Biological Processes (BP), Molecular Functions (MF) and Cellular Compartments (CC). The following parameters were used as cut-offs to represent bubble plots: gene count > 50, fold enrichment > 3 and p-value < 0.01.

#### Interactome analysis

To obtain the protein-protein interaction networks, the 200 more abundant proteins were submitted to STRING (version 11.0). High confidence interactions (minimum required interaction score 0.700) were plotted with hiding of disconnected nodes for ease of representation. Protein clustering was performed using the Markov Cluster Algorithm (MCL) with inflation parameter of 1.4. Highlighted clusters were manually annotated.

#### Venn diagrams

To perform a GO-based analysis of protein content, categories of interest were manually defined from the list of protein hits contained in offspring GO terms of “Extracellular matrix (GO:0031012)”, “Cell-adhesion (GO:0007155)”, “Axon guidance (GO:0007411)” and “Axonogenesis (GO:0007409)”, and from a custom-defined list of glial cells markers (see **Table 2**). For each category, protein hits contained in each brain region were sorted according to their detection in mass spectrometry. For each category, all brain regions were compared using the online tool Venny 2.0.2 (Oliveros, J.C., 2007).

#### PCA analysis

For each brain target, Principal Component Analysis (PCA) was performed to highlight biological differences between injured and intact conditions across replicates. Proteins undetected in more than 5 samples were filtered out. Samples were plotted according to the first and second components, with the percentage of variation indicated for each component.

#### Volcano plots

For each brain target, protein hits were plotted according to the p-value and the log_2_ fold change in injured of differential expression between intact condition. Proteins with a FDR below 5% were highlighted.

#### Heatmaps

To analyse protein expression modulated by the injury, heatmaps were generated by plotting the abundance of each protein hit normalized across all replicates (log_2_ transform). For representation, proteins were selected according their fold change between injured and intact conditions (log_2_ fold change > 0.8) and the FDR-corrected p-value (FDR < 5% for all brain targets, except chiasm: FDR < 1% for ease of representation).

### Transcriptomics datasets data screening

For the screen of gene expression in RGC, GEO datasets available on NCBI were used: atlas of neonatal (P5) RGC from single-cell transcriptomics analysis (accession number GSE115404, (Rheaume et al., 2018)); atlas of adult RGC from single-cell transcriptomics analysis (accession number GSE137400, (Tran et al., 2019)); microarray dataset comparing Pten^-/-^ SOCS3^-/-^ RGC to WT RGC after optic nerve crush (accession number GSE32309, (Sun et al., 2011)); RNA-sequencing dataset comparing Sox11-overexpressing RGC to Plap-overexpressing (control) RGC after optic nerve crush (accession number GSE87046, (Norsworthy et al., 2017)). For single-cell transcriptomics analyses (Rheaume et al., 2018; Tran et al., 2019), online visualization tools were used to determine expression of genes of interest: https://health.uconn.edu/neuroregeneration-lab/rgc-subtypes-gene-browser/ and https://singlecell.broadinstitute.org/single_cell/study/SCP509/mouse-retinal-ganglion-cell-adult-atlas-and-optic-nerve-crush-time-series/. If the mean expression value was strictly positive, the gene was considered to be expressed in the corresponding cluster. For the microarray dataset (Sun et al., 2011), differential gene expression analysis was performed using the interactive web tool GEO2R (NCBI, https://www.ncbi.nlm.nih.gov/geo/info/geo2r.html) to plot the log fold-change and the FDR-corrected p-value. For the RNA-sequencing dataset (Norsworthy et al., 2017), the complete gene list available on NCBI was used to plot the log fold-change and the FDR-corrected p-value.

### Western blot

For each brain taregt, 5-10µg protein were loaded on SDS-PAGE gels (4-15% acrylamide, Biorad). After 1h of electrophoresis at 150V, liquid transfer was performed 2h30 at 320mA on a nitrocellulose membrane (ThermoFisher Scientific). A Ponceau red staining was performed to check the presence of proteins and therefore the quality of protein transfer. Membranes were blocked with 5% milk for 1h at RT and incubated with anti-NCAM1 (1:1000, Rabbit, Cell signalling Technology), anti-TenascinC (1:1000, Rabbit, Abcam), anti-Sema7a (1:1000, Rabbit, Abcam), anti-Nrp2 (1:1000, Rabbit, Cell signalling Technology), anti-PlexinA4 (1:1000, Rabbit, Cell signalling Technology) primary antibodies overnight at 4°C on a shaker. Following incubation with horseradish peroxidase-linked secondary antibody (1:5000, Rabbit, Protein Tech) for 1h at RT, membranes were developed with ECL substrate (10% Tris HCl 1M pH 8.5, 0.5% coumaric acid (Sigma-Aldrich), 0.5% luminol (Sigma-Aldrich) and 0.15% H_2_O_2_ (Sigma-Aldrich)). Data quantification was done using pixel densitometry. For each independent biological replicate, the pixel density of the protein of interest was normalized to actin. Data were normalized to intact condition and subjected to a Student’s t-test for statistical analysis.

### Intracardial perfusion

At the time of sacrifice, mice were anesthesized as described above, then intracardially perfused with ice-cold PBS for 3min and with ice-cold 4% formaldehyde in PBS for 3min. Brains were dissected out and samples were post-fixed overnight at 4°C in 4% formaldehyde.

### Immunofluorescence

Samples were post-fixed in 4% formaldehyde (Sigma) overnight and transferred to 30% sucrose for 2 days at 4°C to cryoprotect. Samples were then embedded in tissue freezing medium compound (MM-France) and frozen at −80°C. 30µm and 20µm thick sections were performed for 10 week brain and young animals, respectively using a cryostat (CryoStar NX50, ThermoFisher Scientific). Immunohistochemistry on tissue sections was performed according to standard procedures. Sections were blocked for 1h with 3% BSA, 5% Donkey Serum, 0.1% PBS-Triton and incubated with the following primary antibodies diluted in the blocking solution, overnight at 4°C: anti-NCAM1 (1:100, Rabbit, Cell signalling Technology), anti-Dclk2 (1:100, Rabbit, Abcam), anti-Sema4D (1:100, Rabbit, Abcam), anti-Col6a3 (1:100, Mouse, Merck Millipore), anti-Cspg4 (1:100, Rabbit, ProteinTech), anti-GFAP (1:200, Rat, ThermoFisher Scientific). Triton was omitted from the blocking solution for NCAM1, Sema4D and L1CAM immunostaining. For L1CAM, CSPG4 and Sema4D, heat-induced antigen retrieval was performed for 5min in citrate buffer. This was followed by incubation with Alexa-fluor conjugated (anti-Rabbit, ThermoFisher Scientific; anti-Mouse, ThermoFisher Scientific; anti-Rat, Jackson Laboratory) antibodies according to standard protocol (dilution 1:200). Slides were mounted with Fluoromount-G with DAPI medium (ThermoFisher Scientific).

### Imaging

Epifluorescence microscope Nikon Ti Eclipse was used for imaging of brain sections.

### Whole-mount tissue clarification and imaging

#### Brain transparization

Whole-brain transparization was performed as described (Renier et al., 2016). For visualization of the brain targets and optic tracts of the visual system, each eye of an adult WT mouse was injected with CTB-555 and CTB-647. After intracardial perfusion, the brain and optic chiasm was dissected out and post-fixed in 4% formaldehyde. The brain was dehydrated in methanol, then bleached overnight in 6% H_2_O_2_ in methanol. After rehydration in PBS, the brain was permeabilized in PBS 0.5% Triton for several days at 4°C. The brain was dehydrated in methanol, then incubated for several hours in dichloromethane/methanol (2:1), then 30min in dichloromethane (Sigma-Aldrich) before transparization in dibenzyl ether (Sigma-Aldrich). Transparized brain was imaged using the lightsheet microscope from LaVision Biotec. Data processing and visualization was performed using Imaris software.

#### Optic nerve clarification

Optic nerve clarification was performed as described (Dodt et al., 2007; Schaeffer et al., 2020). After intarcardial perfusion and post-fixation of the eyes, optic nerves were dissected and dehydrated in ethanol. Optic nerves were incubated for 2hours in hexane, then transparized in benzyl benzoate/benzyl alcohol (2:1) (Sigma-Aldrich). Optic nerves were imaged using a spinning disk confocal microscope (Andor Dragonfly).

## FIGURE LEGENDS

**Table 1: List of proteins detected in each adult visual target**. SCN, suprachiasmatic nucleus; vLGN, ventral lateral geniculate nucleus; dLGN, dorsal lateral geniculate nucleus; SCol, superior colliculus.

**Table 2: List of proteins in each category of interest (ECM, adhesion, axon growth and guidance, glia)**.

**Table 3: List of proteins related to axon guidance and guidance-associated factors in each visual target**. Each tab contains the proteins identified in one visual target and classified according to different categories manually defined (cell adhesion, extracellular matrix, glia components, axon growth and guidance, see **Table 2**). Each entry is given with the protein accession number, gene name and description.

**Table 4: List of protein modulated by the lesion in each adult visual target**. Up-regulated and down-regulated proteins are represented in individual tabs for each visual target.

**Supplementary Figure 1: High reproducibility of the experimental design**. Scatterplots of protein abundance of the detected hits across replicates, in intact (left) and injured (crush, right) conditions. The Pearson’s correlation coefficient is indicated on each plot.

**Supplementary Figure 2: Adult visual targets show similar protein landscape. (A-E)** Network-based cluster analysis of the 200 most abundant proteins identified in each visual target: (A) optic chiasm, (B) SCN, (C) vLGN, (D) dLGN, (E) Superior Colliculus as analyzed with STRING. Only interactions with high confidence (minimum required interaction score > 0.700) are represented. Clustering is done with a Markov Cluster Algorithm (MCL = 1.4). Corresponding functional categories were manually annotated.

**Supplementary Figure 3: Comparison of the vLGN and dLGN and expression of extracellular matrix molecules and glia components in each adult visual target. (A)** Venn diagram showing the rate of similarity (shared detected proteins) between vLGN and dLGN. **(B)** Venn diagrams showing the rate of similarity (shared detected proteins) between vLGN and dLGN in each category as defined in **Table 2. (C)** Venn diagram representing the shared extracellular-matrix-related hits (number of proteins and percentage of the protein number in the extracellular-matrix category). In boxes are indicated proteins uniquelly detected in each visual target. **(D)** Immunofluorescent labelling of CSPG4 expressed in each visual target in an adult intact brain (co-labelled with CTB). Scalebar: 100µm. **(E)** Venn diagram representing the shared glia-related hits (number of proteins and percentage of the protein number in the glia components category). In boxes are indicated proteins uniquelly detected in each visual target. **(F)** Immunofluorescent labelling of GFAP expressed in each visual target in an adult intact brain (co-labelled with CTB). Scalebar: 200µm.

**Supplementary Figure 4: Bilateral optic nerve crush causes modifications of the proteome compared to intact. (A-E)** PCA analysis showing a clustering of the replicates according to the condition intact or injured (crush) in each adult visual target. **(F-H)** Volcano plots showing differentially expressed proteins in the SCN, the dLGN and the SCol, in injured versus intact conditions. Highlighted are protein hits with log_2_ fold change < 0.8 and FDR < 5%. **(I-K)** Heatmaps showing differentially expressed proteins between injured (crush) and intact conditions. The values are abundance normalized across all samples for each protein hit. Proteins with FDR < 5% are represented.

**Supplementary Figure 5: Screen of guidance factors expressed in RGC in intact and injured conditions. (A)** Table representing the expression of guidance receptors corresponding to guidance ligands detected in adult visual targets. In red: repulsive interaction, in green: attractive interaction, in grey: not expressed. For intact atlases (Neonatal RGC (Rheaume et al., 2018) and Adult RGC (Tran et al., 2019)), the number of RGC clusters represented is indicated in the boxes. For regenerative datasets (Pten^-/-^ SOCS3^-/-^ (Sun et al., 2011) and Sox11-OE (Norsworthy et al., 2017)), the variation of the receptor compared to WT (non-regenerative) injured condition is indicated in the boxes if it is upregulated or downregulated (else steady). **(B)** Heatmap showing the variation of expression of guidance or cell adhesion molecules between regenerative models and WT (injured) condition (Pten^-/-^ SOCS3^-/-^ and Sox11-OE. The values are log_2_ fold-change (FC). The genes with significant FC difference (FDR-adjusted p-value < 0.05) are boxed. Sox11-OE: Sox11-overexpressing.

## ETHICS STATEMENT

All animal care and procedures have been approved by the Ethics Committee of Grenoble Institut Neurosciences (project number 201612161701775) and by the French Ministry of Research (project numbers APAFIS#9145-201612161701775v3 and APAFIS#26565-2020061613307385v3) in accordance with French and European guidelines.

## AUTHOR CONTRIBUTIONS

HN and SB conceptualized the study. NV, JS, CD and AP performed experiments. YC and AMH performed mass spectrometry-based proteomic data collection and analysis. NV and JS analyzed and interpreted data. NV and JS drafted figures. NV, JS, SB and HN wrote the manuscript.

## ACKNOWLEDGMENTS

The proteomic experiments were partially supported by Agence Nationale de la Recherche under projects ProFI (Proteomics French Infrastructure, ANR-10-INBS-08) and GRAL, a program from the Chemistry Biology Health (CBH) Graduate School of University Grenoble Alpes (ANR-17-EURE-0003). This work was supported by a grant from ANR to HN (C7H-ANR16C49) and SB (ANR-18-CE16-0007). HN is supported by NRJ Foundation. HN and NV are supported by the European Research Council (ERC-St17-759089). JS is supported by Fondation pour la Recherche Médicale (FRM) postdoctoral fellowship (SPF201909009106).

