## supplemental figures for "Axon guidance modalities in CNS regeneration revealed by quantitative proteomic analysis"

### Supplementary Figure 1

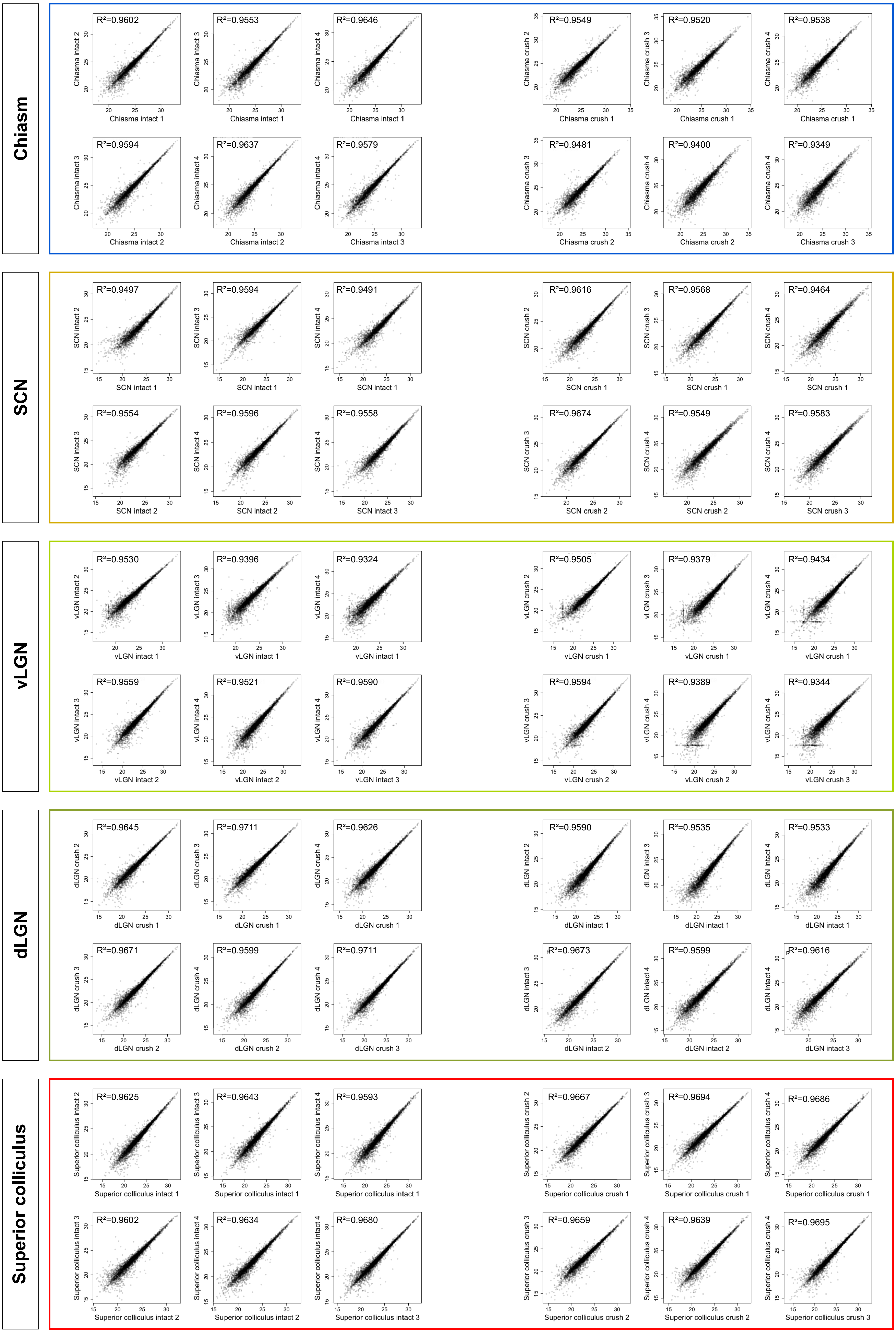

### Supplementary Figure 2

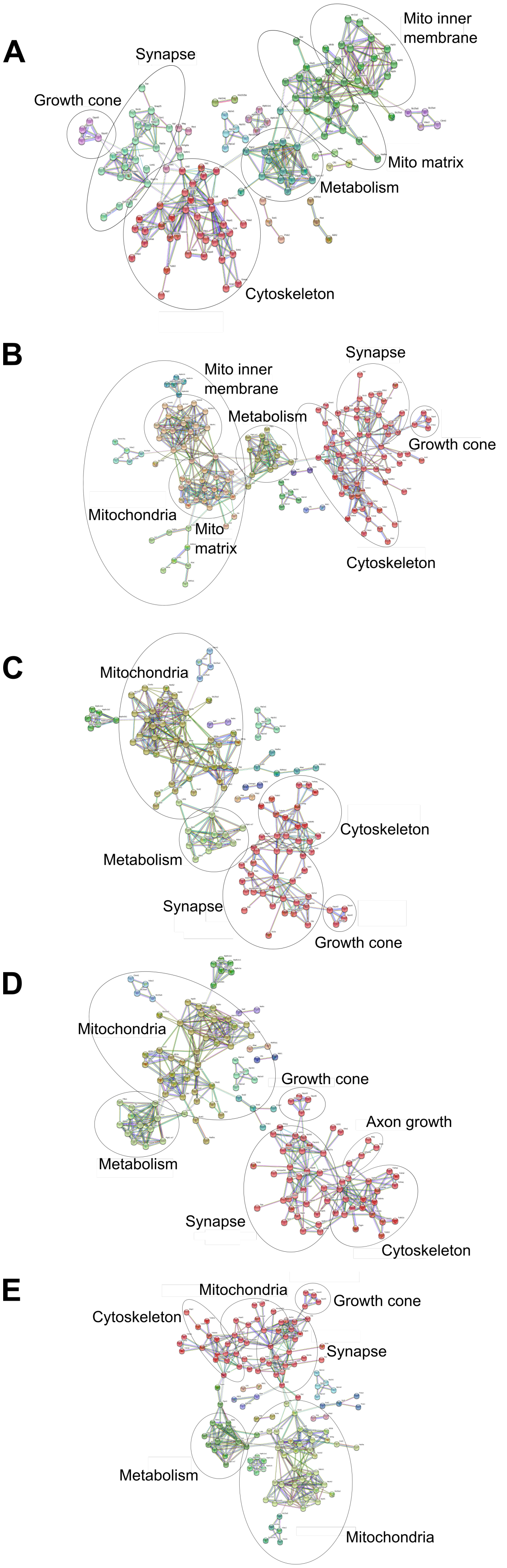

### Supplementary Figure 3

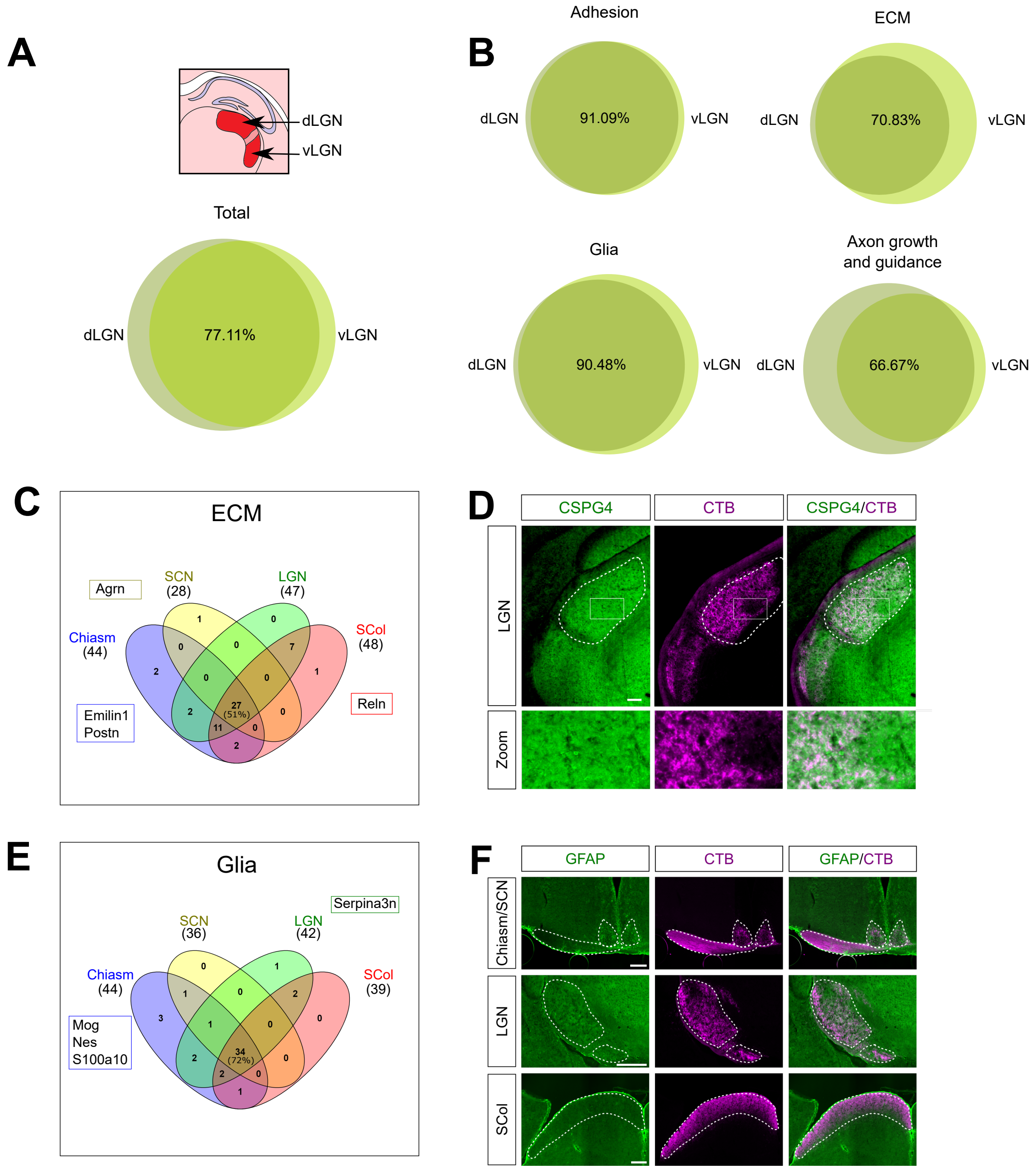

### Supplementary Figure 4

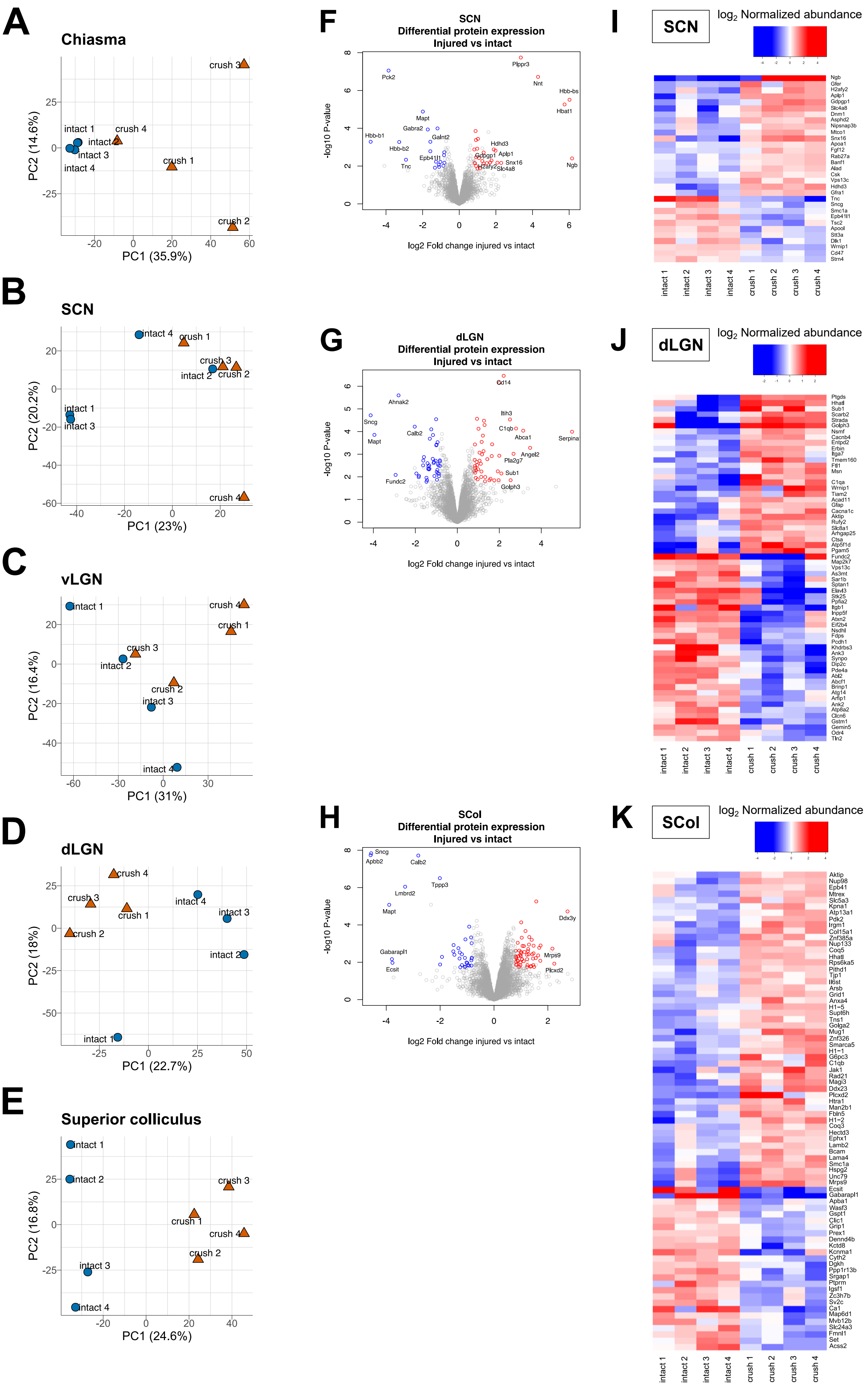

### Supplementary Figure 5

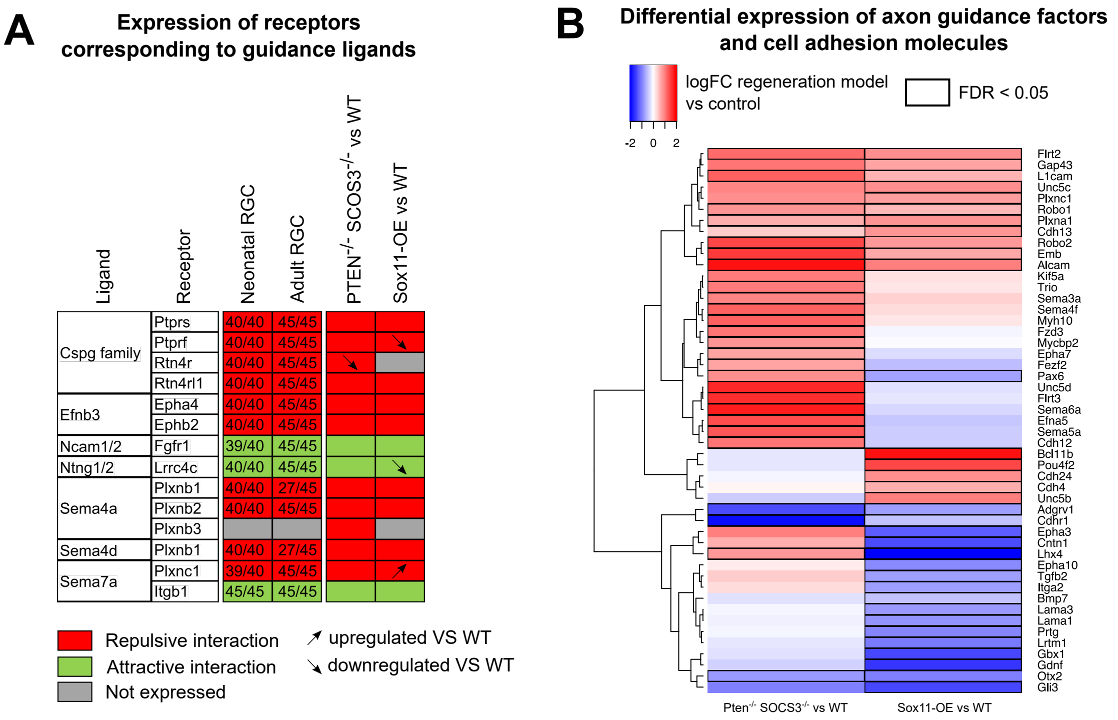
